# Intraspecific trait variation modulates the temperature effect on elemental quotas and stoichiometry in marine *Synechococcus*

**DOI:** 10.1101/2023.09.20.558568

**Authors:** Austin Davis, Nathan S Garcia, Adam C Martiny

## Abstract

Diverse phytoplankton modulate the coupling between the ocean carbon and nutrient cycles through life-history traits such as cell size, elemental quotas, and ratios. Biodiversity is mostly considered at broad functional levels, but major phytoplankton lineages are themselves highly diverse. As an example, *Synechococcus* is found in nearly all ocean regions and contain extensive intraspecific variation. Here, we grew four closely related *Synechococcus* isolates in semi-continuous cultures across a range of temperatures (16-25°C) to quantify for the relative role of intraspecific trait variation vs. environmental change. We report differences in cell size (p<0.01) as a function of strain and clade (p<0.01). The carbon (*Q*_*C*_), nitrogen (*Q*_*N*_), and phosphorus (*Q*_*P*_) cell quotas all increased with cell size. Furthermore, cell size has an inverse relationship to growth rate. Within our experimental design, temperature alone had a weak physiological effect on cell quota and elemental ratios. Instead, we find systemic intraspecific variance of C:N:P, with cell size and N:P having an inverse relationship. Our results suggest a key role for intraspecific life history traits in determining elemental quotas and stoichiometry. Thus, the extensive biodiversity harbored within many lineages may modulate the impact of environmental change on ocean biogeochemical cycles.

## Introduction

Phytoplankton link the ocean carbon, nitrogen, and phosphorus cycles through the biomass ratios of these elements [1–4]. The Redfield ratio describes the ocean C:N:P stoichiometry and is used in models to assess export and productivity [5,6]. However, elemental ratios are variable in surface ecosystem [7,8], and ratios in the interior ocean may change over long-time scales, thereby influencing the ability of the oceans to sequester carbon relative to other nutrient elements [9]. Small cyanobacteria are currently estimated to account for approximately 25% of marine net primary production [10]. Therefore, a clear understanding of how key traits of marine cyanobacteria interact with environmental changes is needed to reduce uncertainties in ocean models of biogeochemistry and net primary production. Elemental ratios within phytoplankton vary with environmental gradients like nutrients [2,11] and phylogenetic origin [12,13]. In addition, previous studies also argue that temperature-dependent mechanisms could influence ratios, but this role is less well established [14,15]

A number of factors are known to influence stoichiometry and elemental quotas. Latitudinal gradients in temperature and community composition are dominant predictors of stoichiometry. However, their respective influences are difficult to decipher and accurately model as they strongly covary in the surface ocean. The nutrient hypothesis predicts that storage of nutrients like P in polyphosphates [16] and N in phycobiliproteins [17,18] may occur under slow growth, but elemental ratios approach an optimum under fast-growth when abundances and activities of components like P-rich ribosomes are high [2,11,19]. Temperature may directly influence the abundance of P-rich ribosomes in cells through a compensatory mechanism for reduced transcriptional activity under low temperature [14]. Variability in sea surface temperature has been documented [20], and future change is projected to cause an increase in sea surface temperatures globally [21]. Cellular elemental stoichiometry is also known to vary between major phylogenetic groups [22]. However, there can also be extensive intraspecific variation in in cell size [23]. Cell size is a master trait that has significant implications for ecology and biogeochemistry, as a number of subordinal traits depend upon this key trait [24]. The dominance of small microbes in the open ocean facilitates rapid cycling of carbon by the microbial loop [25]. Sinking velocity increases with cell size, and aggregates of cells may form larger particles, which may more rapidly sink and lead to greater carbon sequestration [26]. The metabolic rate has been reported to scale inversely with cell size, with the degree of variance dependent on phylogeny [26–30]. Nutrient diffusion and uptake are enhanced in smaller cells due to their greater surface area to volume ratio [26,31]. Thus, cell size may influence elemental stoichiometry through these subordinal traits. In the open ocean, cell size varies inversely with population abundance [32], thereby exerting strong influences on ecology and biogeochemical models that rely on abundance [6,10,33–35]. If temperature deviates from the optimal range (T_opt_), variation in cell size may occur [26]. Key factors such as temperature and nutrient availability are considered major factors in determining certain biogeochemical estimates, such as net primary production, but cell size must also be considered [10].

At the genus level, *Synechococcus* is one of the most productive lineages in the world and plays an important role at the base of oceanic food webs along broad nutrient and thermal gradients [10]. Within *Synechococcus*, multiple clades have been cited as being differently thermally adapted, with some clades more dominant in cold, nutrient-rich water relative to others [36,37]. Variable thermal optima for growth may be key in contributing to differences in biogeographical dominance of clades [38] but unknown traits may influence its biogeography. Variability in cell size among lineages of *Synechococcus* is known to exist [2,38] but has not been thoroughly examined despite the roles that cell size may play in determining cellular growth rate, elemental quotas, and stoichiometry.

Here, we ask the following questions: 1) How does cell size vary between lineages of *Synechococcus*? 2) Are elemental quotas and ratios systematically different between strains of *Synechococcus*? 3) How does temperature influence growth rate, elemental quotas, and ratios of *Synechococcus*? We hypothesized cell size, stoichiometry, and elemental quotas would all be closely and strongly linked to *Synechococcus* strain identity. We also hypothesized strong thermal effects on stoichiometry, elemental quotas, and growth rate, as per the translation-compensation hypothesis, with a greater amount of phosphorus found in cells with elevated growth rates or under lower experimental temperatures. To test these hypotheses, we examined four *Synechococcus* strains, representing a gradient in cell size, from two cold-adapted clades isolated from three locations across the world: CC9902, BL107 (clade IV), CC9311, and ROS8604 (clade I) [39–42]. We present new findings about the potential link between cell size and phylogeny of *Synechococcus*, raising new questions about the ecology and biogeochemistry of picocyanobacteria.

## Materials and Methods

We incubated semi-continuous cultures of *Synechococcus* (CC9311, BL107, ROS8604, CC9902, representing two clades) (Fig 1, Table 1) in triplicate 1 L flasks at 16°C, 18°C, 20°C, 22°C, 25°C, and 27°C. Ambient light (60 μmol quanta m^-2^ s^-1^) was supplied using white fluorescent lamps on a 12:12 light-dark cycle. Culture media (modified artificial sea water) was prepared as described in Garcia et al. (2016) [2]. We supplied nitrate (NO_3_^-^) and phosphate (PO_4_^3−^) in concentrations of 125 μM and 10 μM, respectively. We transferred media and diluted cultures using an open flame in a hood in order to avoid contamination. We routinely diluted to maintain a stable growth rate.

**Table 1:** *Synechococcus* strain information. The strains used during experimentation are presented here, with additional information such as clade, location of isolation, previously reported temperature optimum, and range in cell size, median cell size, and mean cell size gathered during the course of experimentation.

| Strain | Clade | Location | Isolation<br>depth (m) | T<sub>Opt</sub><br>(reported) | T<sub>Opt</sub><br>(observed) | Cell size<br>(FSCH) range<br>(sampling) | Cell size<br>(FSCH) mean<br>(sampling) | Cell<br>diameter<br>(μm) range | Cell diameter<br>(μm) mean<br>(sampling) |
| --- | --- | --- | --- | --- | --- | --- | --- | --- | --- |
| CC9311 | I [36] | California<br>Current [44] | 95 m | 22°C [45] | 25°C | 33612-105680 | 57968 | 1.50-1.30 | 1.15 |
| BL107 | IV<br>[36,37] | Balearic<br>Sea [40] | 1800 m | 25°C [46] | 22°C | 16126-20826 | 18999 | 0.90-0.95 | 0.93 |
| ROS8604 | I<br>[36,37] | English<br>Channel<br>[42] | 0 m | N/A | 18°C | 65927–88051 | 74855 | 1.20-1.30 | 1.22 |
| CC9902 | IV<br>[36,37] | California<br>Current [39] | 5 m | N/A | 16°C | 4639–8021 | 5636 | 0.64-0.75 | 0.67 |

**Fig 1.**
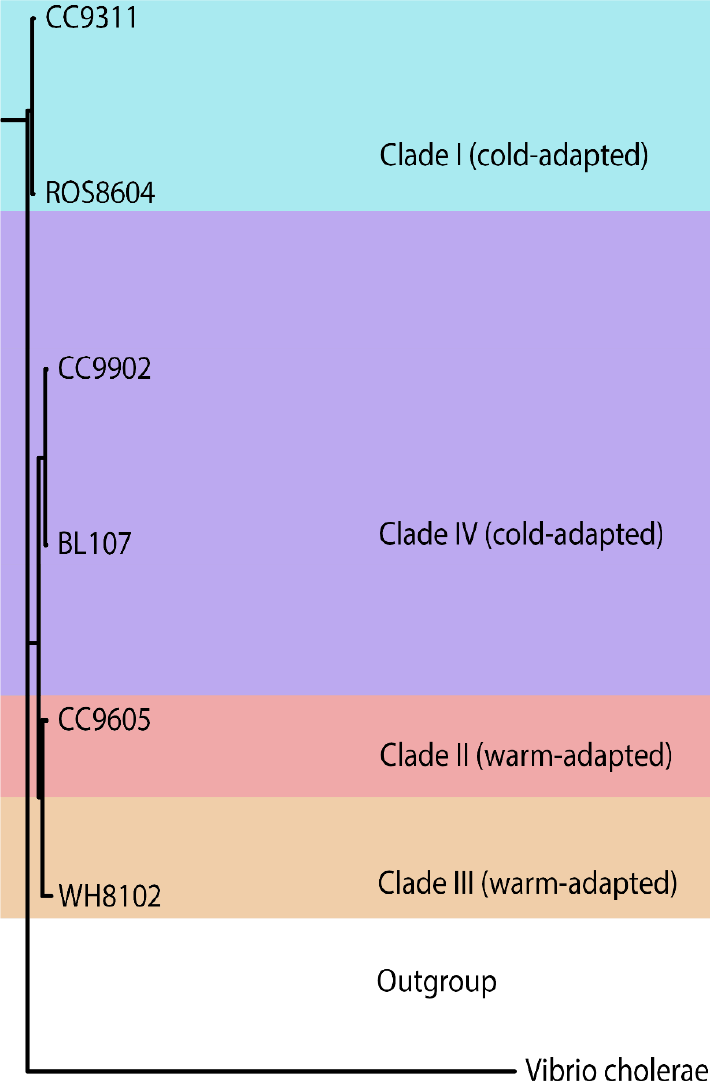
Phylogeny of examined *Synechococcus* strains. Color represents strain plotted. *Vibrio cholerae* serves as the outgroup for constructed *Synechococcus* phylogeny. CC9605 and WH8102 represent clades II and III, respectively.

We measured particulate organic carbon (POC), particulate organic nitrogen (PON), and particulate organic phosphorus (POP), as well as cell enumeration by flow cytometry, following the methods outlined in Garcia et al. (2016) [2]. We sampled after seven doublings or one month to acclimate cells to the temperature conditions. We vacuum filtered POC, PON (150 mL), and POP (50 mL) samples onto pre-combusted GF/F Whatman glass filters (450°C) at 10 psi. We dried particulate organic carbon and particulate organic nitrogen samples at 50-80°C for a minimum of 48 hours and pelletized prior to analysis using a Flash EA 1112 NC Soil Analyzer (Thermo-Scientific). We rinsed particulate organic phosphorus samples with 0.17 M NaSO_4_, immersed in 2 mL of MgSO_4_, dried at 80°C overnight, and combusted at 450°C for 2 hours. We then added 5 mL of 0.2 M HCl and baked the samples at 80-90°C. We measured particulate organic phosphorus samples via colorimetric assay following the Bermuda Atlantic Time-series methodology [46] using a Genesys 10S UV-vis spectrophotometer (Thermo-Scientific) at 885 nm.

Culture cell density and growth (cell growth greater than that of cell death) rate was measured every two-three days and immediately prior to sampling using a NovoCyte 1000 flow cytometer (excitation laser 488 nm, emission peak 575 nm) and forward scatter. We counted the cells at a flow rate of 35 μL/min. To assess heterotrophic populations, we stained the cultures with SYBR Green (Thermo-Fisher) for 15 minutes at room temperature, vortexed them, and counted using the FITC (excitation laser 488 nm, emission peak 520 nm) channel. We recorded duplicate cell counts at each sampling.

We measured cell diameter by microscopy under oil immersion at 1000x magnification using the Axioplan2 and AxioView 1.4.5 sizing software (Carl Zeiss, Goettingen, Germany) with reference to a staged micrometer (Ted Pella Inc., Redding, CA). To estimate cell diameter, we created a conversion factor determined by plotting the mean observed cell diameters and mean forward scatter values for several strains of *Synechococcus* (Fig 2).

**Fig 2.**
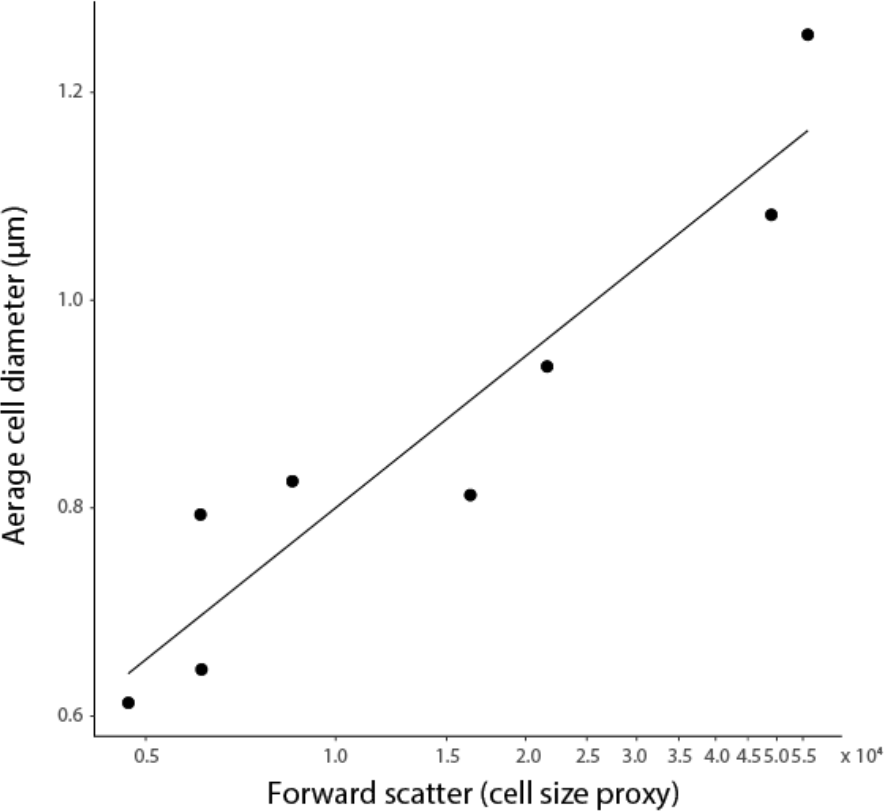
Cell diameter is linked to forward scatter. Cell size measured using flow cytometry (forward scatter, FSCH) and microscopy are highly correlated (r^2^=0.83, p=0.001).

We used the R statistical software (www.r-project.org) to perform linear discriminant analyses, analysis of variance, and multivariate analysis of variance analyses. We used the package PHYLIP (https://evolution.genetics.washington.edu/phylip.html) to construct a phylogenetic tree using *rpoc1* sequences (BioCyc, NCBI).

## Results

We sought to assess the role of intraspecific life history traits vs. temperature on cell size, elemental quotas, growth rate and stoichiometry in *Synechococcus*, a widely distributed key producer and highly ubiquitous picophytoplankton (Table 1). We grew semicontinuous cultures and recorded the growth rate every other day for the strains ROS8604, BL107, CC9902, and CC9311. We grew cultures at 16°C, 18°C, 20°C, 22°C, 25°C, and 27°C and subjected them to a 12-hour light/12-hour dark cycle. We performed cell counts and assessment of cell size (forward scatter, FSCH) every other day using flow cytometry. Cell diameter was calculated from paired FSCH and microscopy measurements of *Synechococcus* (Fig. 1). We sampled for cellular carbon, nitrogen, and phosphorus quotas after an acclimation period of at least seven doublings or three weeks (for strains with slower growth rates). As we did not observe growth which was greater than cell death at 27°C, we conclude this is the upper thermal limit for each of the four strains under our experimental conditions.

We observed distinct ranges in cell size linked to strain identity and phylogeny, whereas temperature had weak effects. We find hierarchical effects on *Synechococcus* cell size. The greatest influence on cell size is clade. Clade I strains ROS8604 and CC9311 exhibited the largest cell sizes, while Clade IV strains BL107 and CC9902 were substantially lower in cell size (Table 1, Fig 3). Inter-clade differences in the effects on cell size were greater than intra-clade effects (Fig 3). However, our results indicated strain served as a secondary phylogenetic effect on cell size (Table S2). In contrast to the effects of clade and strain on cell size, we observed weaker thermal effects on cell (Table S1). As such, temperature only had a significant effect on cell size when accounting for intraspecific strain variations (Table S1).

**Fig 3.**
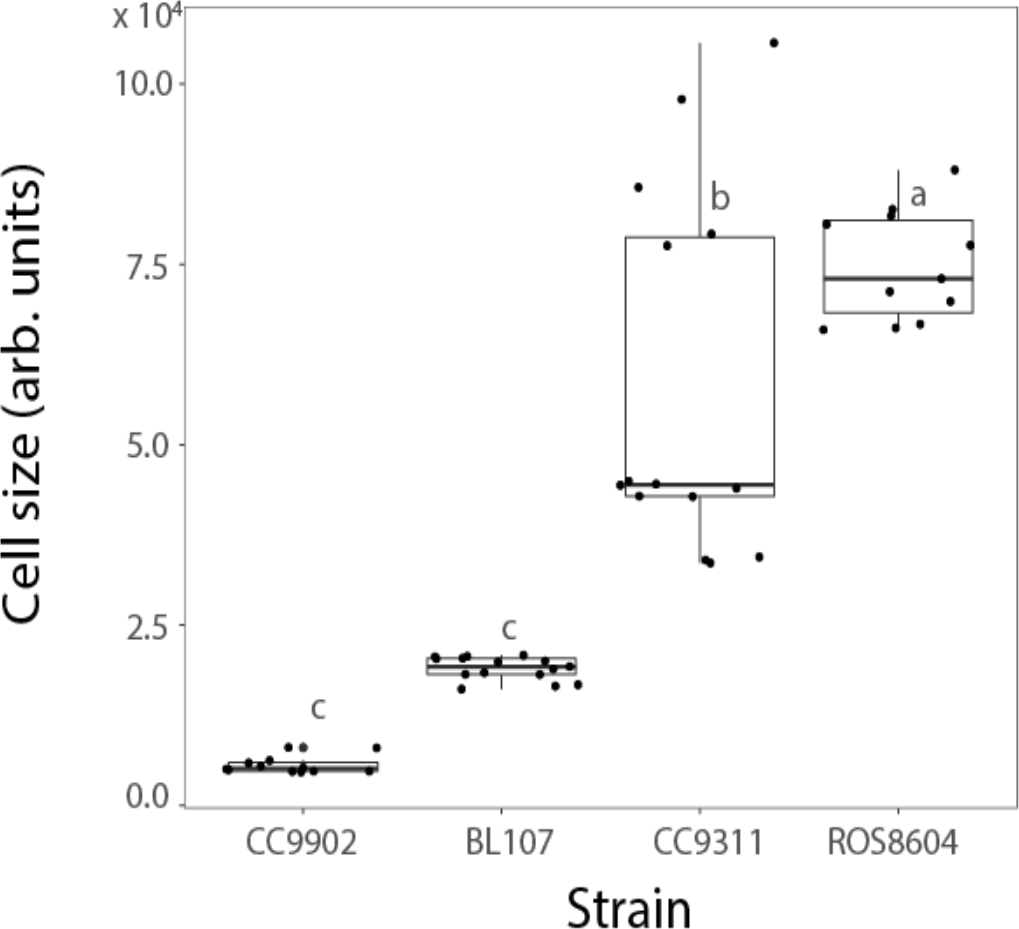
Intraspecific cell size variation among *Synechococcus*. Ranges in cell size (FSCH) across strains of *Synechococcus*. We compared the effects of strain on cell size using a one-way ANOVA and Tukey’s honest significance difference (HSD) test, represented by compact letter display (CLD).

We next investigated the relative role of intraspecific trait variation vs. temperature on elemental quotas. The cellular carbon quota (*Q*_*C*_) displayed a strong positive relationship with cell size (Fig 4A). A similar pattern was also seen between cell size and the nitrogen quota (*Q*_*N*_) and the phosphorus quota (*Q*_*P*_) (Fig 4). As cell sizes were unique among the strains, significant relationships between all cell quotas and strain identity were seen (Table S2). However, we did not find effects of thermal effects alone on *Q*_*P*_, *Q*_*C*_ or *Q*_*N*_, but did observe the presence of thermal effects when other factors, such as strain, were considered for each of these elemental quotas (Table S4, Table S5). *Synechococcus* represents a diversity of elemental quotas within a limited selection of strains; *Q*_*C*_ ranges from 35.4 fg to 520 fg, *Q*_*N*_ ranges from 10.0 fg to 96.7 fg, and Q_*P*_ ranges from 1.23 fg to 14.7 fg. The *Synechococcus* strains we studied exhibit elemental quota ranges of 14.7 times for *Q*_*C*_, 9.7 for *Q*_*N*_, and 11.4 for *Q*_*P*_ (Fig 4). The degree of variability in elemental quotas is dependent upon lineage; for example, Clade I strains exhibit a wider range in elemental quotas than Clade IV strains (Fig 4A, Fig 4C).

**Fig 4.**
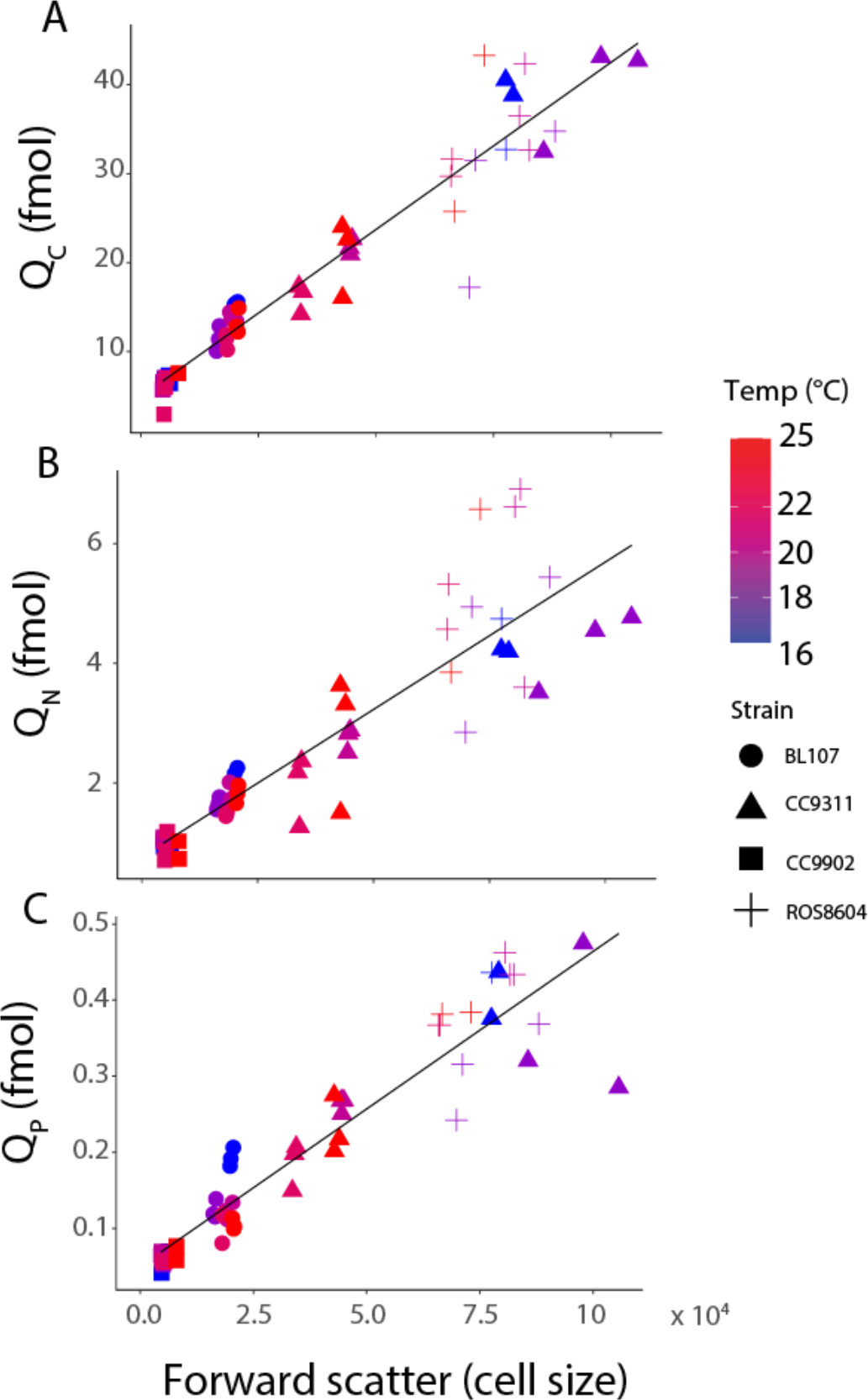
Elemental quotas scale with cell size across *Synechococcus* strains. Fig 4A depicts the carbon cell quota (*Q*_C_) with forward scatter, Fig 4B depicts nitrogen cell quota *Q*_*N*_ with forward scatter, and Fig 4C depicts *Q*_*P*_ with forward scatter. Point shape represents strain plotted. Point color represents temperature. We performed principal component regressions for each elemental quota and cell size.

The intraspecific differences in cell size influenced stoichiometry, while the degree and precise nature of the influence varied (Fig 5). We observed significant relationships between strain and both the C:N and N:P ratios (Fig 5B, Table S2). In contrast, we found no relationship between strain and C:P ratio (Fig 5C, Table S2). However, when stoichiometric response was examined by clade, we did observe significant differences in N:P and C:P (Table S3; Fig S3) but not for C:N ratio (Table S3; Fig S3). We found no effects of temperature alone on the N:P, C:N, or C:P (Table S1). While we did not observe a significant relationship between cell size and C:N or C:P, we did determine a highly significant, inverse relationship between cell size and N:P (Fig 5D-F, Table S6). We observed the greatest range in C:N for CC9902 (4.10-10.0), and the lowest range in C:N in BL107 (6.5-8.0). We found CC9902 was the strain with the greatest range in N:P (9.6-22.0), with the smallest range found in BL107 (11-19). We found C:N mean values were 6.30 (CC9902), 7.30 (BL107), 8.60 (CC9311), and 6.60 (ROS8604). We observed the lowest C:P mean (ROS8604, 113.0) for the largest strain examined (CC9902, 155.0; BL107, 145.0; CC9311; 150.0).

**Fig 5.**
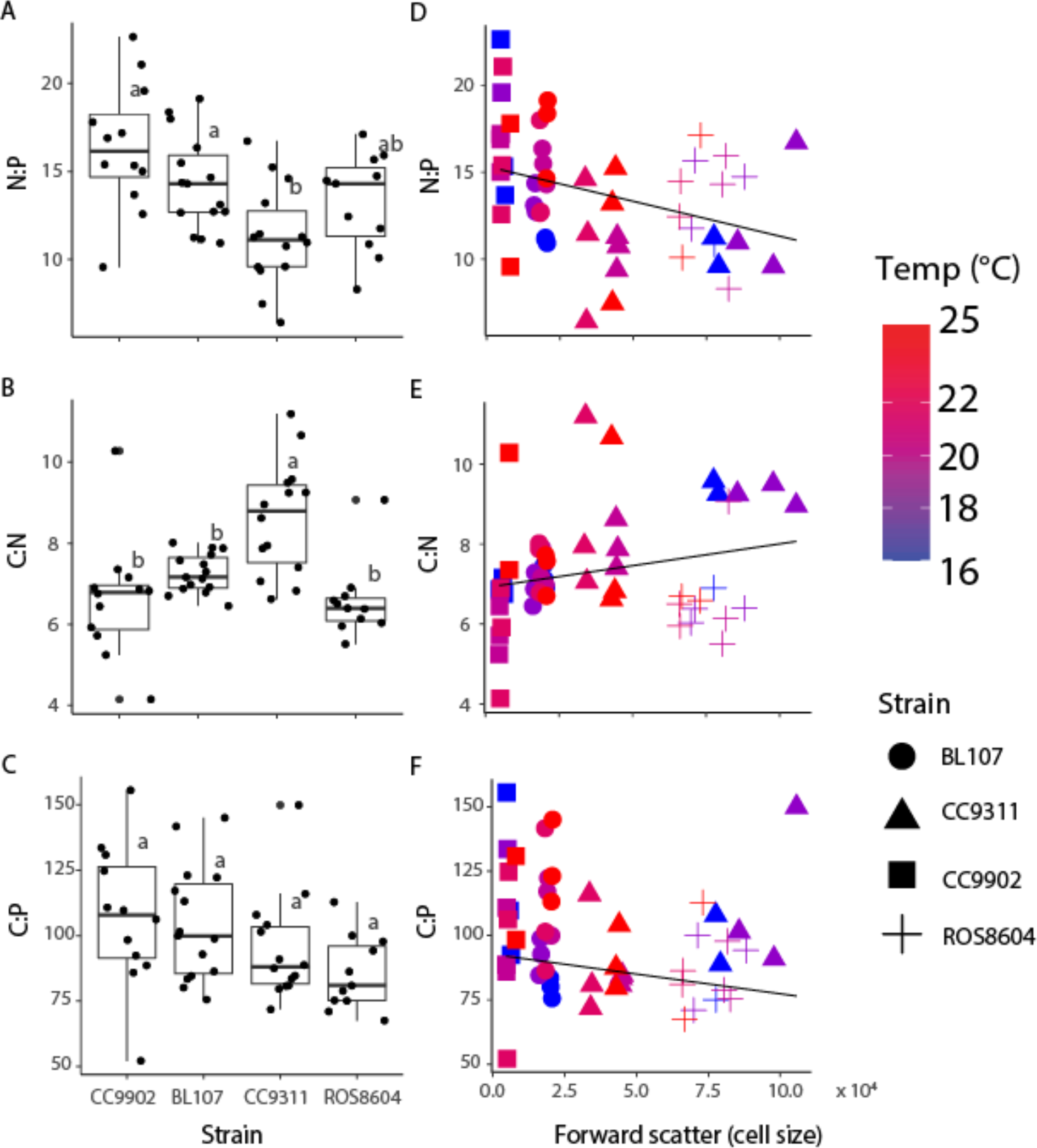
Interactions between *Synechococcus* strain and stoichiometry, and cell size and stoichiometry. We depict effects of strain on N:P (Fig 5A), C:N (Fig 5B), and C:P (Fig 5C) stoichiometry. We depict effects of cell size on N:P (Fig 5D), C:N (Fig 5E), and C:P (Fig 5F) stoichiometry. Color represents temperature (Fig 5A-C). We compared the effects of strain on stoichiometry (Fig 5A-C) using a one-way ANOVA and Tukey’s honest significance difference (HSD) test, and are represented by compact letter display (CLD). Elemental ratios are molar.

We next detected a link between cell size, growth rate and elemental quotas, with a weaker influence of temperature. To test the growth rate hypothesis for *Synechococcus* strains, we assessed the cellular phosphorus quota (Q_P_) and growth rate at different temperatures in Clade IV and I. We observed an inverse relationship between growth rate and Q_P_ (Fig 6A, Table S6) and thus the opposite of this hypothesis. Furthermore, growth rate was influenced by temperature only when strain was also considered as a factor likely reflecting the additional effect of adaptation (Fig 6D, Table S4). Cell size and growth rate varied as a function of both clade and strain, with the two smaller strains (clade IV) demonstrating a greater growth rate than the two larger strains (clade IV) (Fig 6E-G, Table S2, Table S3).

**Fig 6.**
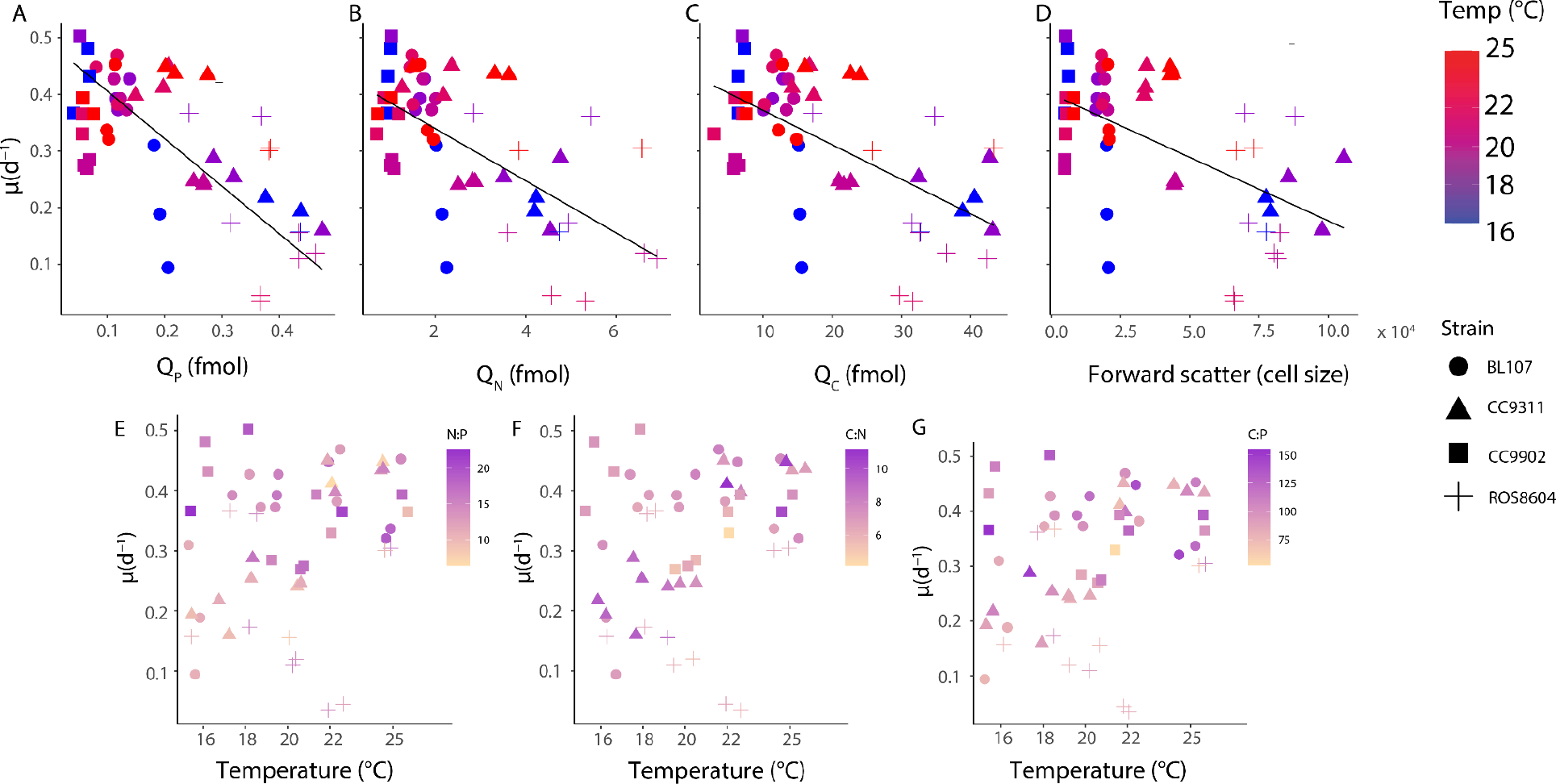
Influence of growth rate on elemental quotas, cell size, and stoichiometry. Growth rate and elemental quotas Q_P_ (Fig 6A), Q_N_ (Fig 6B), and Q_C_ (Fig 6C), cell size (Fig 6D), N:P (Fig 6E), C:N (Fig 6F), and C:P (Fig 6G). Color represents temperature (Fig 6A-D) or stoichiometry (Fig 5E-F). Point shape represents strain. We performed principal component analyses of *μ* and Q_P_ (Fig 6A), Q_N_ (Fig 6BF), Q_C_ (Fig 6C) and cell size (Fig 6D), represented by the regression lines. Elemental ratios are molar.

## Discussion

We observed substantial intraspecific differences in stoichiometric responses among *Synechococcus* strains. We reported higher C:N values relative to previous research conducted using the warm-adapted clade IIIa (WH8012 and WH8103) [36,43]. We found lower C:P ranges in clades I and IV (52-155) relative to those reported for clade IIIa (121-165) [43]. However, while the collective N:P ratios varied between phyla, the grand mean of N:P ratios across all strains of our nutrient-replete *Synechococcus* during exponential growth (13.7) aligned with previously reported values for WH8103 (clade IIIa) (15) and WH7803 (clade V) (13.3) [43,44]. This alignment was only observed during the exponential growth phase (in which we collected samples) emphasizing the importance of growth stage [15,26,45]. Additionally, strains across clade I and IV consistently deviated from the Redfield proportions, supporting the idea that isolates differ from the average ratio of 106:16:1 [5,18].

Our estimates of cell size clearly partitioned isolates into distinct size classes within the commonly identified range for *Synechococcus* [35]. Our estimates for cell diameter were considerably smaller than some previous estimates [46]. As predicted from the metabolic theory of ecology, cell diameter was inversely related to growth rate [17,47], perhaps due to differences in surface area to volume ratios [26]. When combined, low nutrient quotas and high growth rates enable small cells to reach high abundances, particularly in nutrient-poor waters [48]. However, the advantages of being small are counteracted by the costs associated with grazing pressure, which accelerates the trophic transfer of carbon through ecosystems. Conversely, large cells have a higher sinking velocity and relative contribution to carbon export [26]. Modeled estimates of net primary production (NPP), which rely on estimates of *Q*_*C*_ (NPP = *μ* x *Q*_*C*_ x N_cell_) and are typically applied to broad phytoplankton groups [6,34]; as differences in cell size may affect differences in carbon exported, our reported variability in cell size within *Synechococcus*—and the associated linkage with *Q*_*C*_ — highlight the importance of considering strain related variability in *Q*_*C*_. Our results align with previous literature establishing variability in cell size within related phytoplankton groups [49–51].

Contrasting our hypothesis that we would observe strong direct thermal effects, we found temperature exerted little to no direct effect on cell quotas and stoichiometry under nutrient replete semi-continuous growth. Strain-specific temperature responses have been reported in *Prochlorococcus* [15] but the lack of response observed in a previous study on *Synechococcus* was proposed to result from studying *Synechococcus* exclusively at the genus level, and thus only dominant taxa were assessed [52]. As temperature is believed to influence the biodiversity and distribution of phytoplankton, and the optimal C:N:P content is known to vary based on taxonomy or specific oceanic regions [8,22], we also questioned how C:N:P varies across isolates. Although temperature is thought to affect the C:N:P ratios of phytoplankton through the translation-compensation hypothesis [14], we did not find this effect in our assessment of multiple strains of *Synechococcus* across two clades from three distinct regions. We report the observation of indirect thermal effects (e.g., thermal effects as a function of strain). A possible explanation for the lack of strong direct thermal effects may be the lack of our experimental assessment of the lower thermal limits of the four *Synechococcus* strains. Thus, we are unable to make statements about the lower temperature ranges (i.e., approaching thermal limits), where *Synechococcus* may be capable of growth under significant thermal stress. However, we grew *Synechococcus* within a broad temperature range of 16–27°C, with 27°C being the upper thermal limit under our experimental conditions. Another explanation is tied to the semi-continuous method of culturing *Synechococcus* cells. Here, nutrient levels are elevated possibly resulting in nutrient storage, which may have obscured the role that temperature may play in determining physiological response cell quotas and stoichiometry. Thus, a temperature effect may be more pronounced in polar regions or in conjunction with severe nutrient limitation where nutrient storage is depleted.

Phytoplankton play an essential role in biogeochemical cycling through the ratios of carbon, nitrogen, and phosphorus inside cells. It is crucial to understand the environmental effects on picocyanobacterial traits in order to provide more accurate projections of net primary production (NPP) and their contribution to biogeochemical cycling. We provide evidence that intraspecific variation may play a role in the stoichiometric response and cell size of *Synechococcus*. Our findings of distinct size classes within *Synechococcus* evoke those of other phytoplankton phyla, and thus underscore the importance of considering intraspecific trait variation in biogeochemical and productivity models to generate accurate projections of future changes.

## Conflict of Interest

The authors declare no conflict of interest.

## Acknowledgements

Brian Palenik (UC San Diego) provided cultures used in our experimentation. Claudia Weihe and Dr. Jennifer Martiny permitted use of the AxioPlan 2 scope and AxioView imaging software employed in the conversion of FSCH to cell diameter. This work was supported by the National Science Foundation OCE-2135035 and IOS-2137339 to A.C.M. and N.S.G.

## Notes

### Competing Interest Statement

The authors have declared no competing interest.

## References

1. Falkowski PG. Evolution of the nitrogen cycle and its influence on the biological sequestration of CO_2_in the ocean. Nature 1997;387: 272–275. doi:10.1038/387272a0

2. Garcia NS, Bonachela JA, Martiny AC. Interactions between growth-dependent changes in cell size, nutrient supply and cellular elemental stoichiometry of marine Synechococcus. ISME J. 2016;10: 43–69. doi:10.1038/ISMEJ.2016.50

3. Moreno AR, Martiny AC. Ecological stoichiometry of ocean plankton. 101146/annurev-marine-121916-063126. 2018;10: 43–69. doi:10.1146/ANNUREV-MARINE-121916-063126

4. Zehr JP, Kudela RM. Nitrogen cycle of the open ocean: from genes to ecosystems. 101146/annurev-marine-120709-142819. 2010;3: 197–225. doi:10.1146/ANNUREV-MARINE-120709-142819

5. Redfield AC. The biological control of chemical factors in the environment. Sci Prog. 1960;11:150–170.

6. Moore JK, Doney SC, Lindsay K. Upper ocean ecosystem dynamics and iron cycling in a global three-dimensional model. Global Biogeochem Cycles. 2004;18: 1–21. doi:10.1029/2004GB002220

7. Martiny AC, Pham CTA, Primeau FW, Vrugt JA, Moore JK, Levin SA, et al. Strong latitudinal patterns in the elemental ratios of marine plankton and organic matter. Nature Geosci 2013 64. 2013;6: 279–283. doi:10.1038/ngeo1757

8. Tanioka T, Garcia CA, Larkin AA, Garcia NS, Fagan AJ, Martiny AC. Global patterns and predictors of C:N:P in marine ecosystems. Commun Earth Environ. 2022;3. doi:10.1038/S43247-022-00603-6

9. Walker AP, Zaehle S, Medlyn BE, De Kauwe MG, Asao S, Hickler T, et al. Predicting long-term carbon sequestration in response to CO_2_enrichment: How and why do current ecosystem models differ? Global Biogeochem Cycles. 2015;29: 476–495. doi:10.1002/2014GB004995

10. Flombaum P, Gallegos JL, Gordillo RA, Rincón J, Zabala LL, Jiao N, et al. Present and future global distributions of the marine Cyanobacteria Prochlorococcus and Synechococcus. Proc Natl Acad Sci U S A. 2013;110: 9824–9829. doi:10.1073/PNAS.1307701110

11. Rhee G -Y. A continuous culture study of phosphate uptake, growth rate and polyphosphate in Scenedesmus sp. J Phycol. 1973;9: 495–506. doi:10.1111/J.1529-8817.1973.TB04126.X

12. Ho TY, Quigg A, Finkel Z V., Milligan AJ, Wyman K, Falkowski PG, et al. The elemental composition of some marine phytoplankton. J Phycol. 2003;39: 1145–1159. doi:10.1111/J.0022-3646.2003.03-090.X

13. Quigg A, Irwin AJ, Finkel Z V. Evolutionary inheritance of elemental stoichiometry in phytoplankton. Proceedings Biol Sci. 2011;278: 526–534. doi:10.1098/RSPB.2010.1356

14. Toseland A, Daines SJ, Clark JR, Kirkham A, Strauss J, Uhlig C, et al. The impact of temperature on marine phytoplankton resource allocation and metabolism. Nature Clim Chang 2013 311. 2013;3: 979–984. doi:10.1038/nclimate1989

15. Martiny AC, Ma L, Mouginot C, Chandler JW, Zinser ER. Interactions between thermal acclimation, growth rate, and phylogeny influence Prochlorococcus elemental stoichiometry. PLoS One. 2016;11. doi:10.1371/JOURNAL.PONE.0168291

16. Rao NN, Kornberg A. Inorganic polyphosphate supports resistance and survival of stationary-phase Escherichia coli. J Bacteriol. 1996;178: 1394–1400. doi:10.1128/JB.178.5.1394-1400.1996

17. Geider R, Piatt T, Raven J. Size dependence of growth and photosynthesis in diatoms: a synthesis. Mar Ecol Prog Ser. 1986;30: 93–104. doi:10.3354/MEPS030093

18. Geider RJ, La Roche J. Redfield revisited: variability of C:N:P in marine microalgae and its biochemical basis. 2002;37: 1–17. doi:10.1017/S0967026201003456

19. Klausmeier CA, Litchman E, Daufresne T, Levin SA. Phytoplankton stoichiometry. Ecol Res. 2008;23: 479–485. doi:10.1007/S11284-008-0470-8

20. Metz B, Meyer L, Bosch P. Climate Change 2007 - Mitigation of Climate Change. Cambridge University Press;2007. doi:10.1017/CBO9780511546013

21. Amos CL, Al-Rashidi TB, Rakha K, El-Gamily H, Nicholls RJ. Sea surface temperature trends in the coastal ocean. 2013.

22. Quigg A, Finkel Z V., Irwin AJ, Rosenthal Y, Ho T-Y, Reinfelder JR, et al. The evolutionary inheritance of elemental stoichiometry in marine phytoplankton. Nature 2003 4256955. 2003;425: 291–294. doi:10.1038/nature01953

23. Garcia NS, Yung CM, Davis KM, Rynearson T, Hunt DE. Draft genome sequences of three bacterial isolates from cultures of the marine diatom Thalassiosira rotula. Genome Announc. 2017;5. doi:10.1128/GENOMEA.00316-17

24. Andersen KH, Aksnes DL, Berge T, Fiksen Ø, Visser A. Modelling emergent trophic strategies in plankton. J Plankton Res. 2015;37: 862–868. doi:10.1093/PLANKT/FBV054

25. Azam F, Fenchel T, Field JG, Gray J, Meyer-Reil L, Thingstad F. The ecological role of water-column microbes in the sea. Mar Ecol Prog Ser. 1983;10: 257–263. doi:10.3354/MEPS010257

26. Finkel Z V., Beardall J, Flynn KJ, Quigg A, Rees TA V., Raven JA. Phytoplankton in a changing world: cell size and elemental stoichiometry. J Plankton Res. 2010;32: 119–137. doi:10.1093/PLANKT/FBP098

27. Marañón E. Cell size as a key determinant of phytoplankton metabolism and community structure. http://dx.doi.org/101146/annurev-marine-010814-015955. 2015;7: p241–264. doi:10.1146/ANNUREV-MARINE-010814-015955

28. Moloney CL, Field JG. General allometric equations for rates of nutrient uptake, ingestion, and respiration in plankton organisms. Limnol Oceanogr. 1989;34: 1290–1299. doi:10.4319/LO.1989.34.7.1290

29. Raven J, Finkel Z, Irwin A D.D. D. Picophytoplankton : Bottom-up and top-down controls on ecology and evolution. 2005.

30. Teng YC, Primeau FW, Moore JK, Lomas MW, Martiny AC. Global-scale variations of the ratios of carbon to phosphorus in exported marine organic matter. Nature Geosci 2014 712. 2014;7: 895–898. doi:10.1038/ngeo2303

31. Stolte W, Riegman R. Effect of phytoplankton cell size on transient-state nitrate and ammonium uptake kinetics. Microbiology. 1995;141: 1221–1229. doi:10.1099/13500872-141-5-1221

32. Agusti S, Duarte CM, Kalff J. Algal cell size and the maximum density and biomass of phytoplankton. Limnol Oceanogr. 1987;32: 983–986. doi:10.4319/LO.1987.32.4.0983

33. Behrenfeld MJ, Boss E, Siegel DA, Shea DM, Behrenfeld MJ, Boss E, et al. Carbon-based ocean productivity and phytoplankton physiology from space. Global Biogeochem Cycles. 2005;19: 1–14. doi:10.1029/2004GB002299

34. Edwards KF, Thomas MK, Klausmeier CA, Litchman E. Light and growth in marine phytoplankton: allometric, taxonomic, and environmental variation. Limnol Oceanogr. 2015;60: 540–552. doi:10.1002/LNO.10033/SUPPINFO

35. Westberry T, Behrenfeld MJ, Siegel DA, Boss E. Carbon-based primary productivity modeling with vertically resolved photoacclimation. Global Biogeochem Cycles. 2008;22: 2024. doi:10.1029/2007GB003078

36. Sohm JA, Ahlgren NA, Thomson ZJ, Williams C, Moffett JW, Saito MA, et al. Cooccurring Synechococcus ecotypes occupy four major oceanic regimes defined by temperature, macronutrients and iron. ISME J. 2016;10: 333–345. doi:10.1038/ISMEJ.2015.115

37. Stuart RK, Brahamsha B, Busby K, Palenik B. Genomic island genes in a coastal marine Synechococcus strain confer enhanced tolerance to copper and oxidative stress. ISME J 2013 76. 2013;7: 1139–1149. doi:10.1038/ismej.2012.175

38. Lopez JS, Garcia NS, Talmy D, Martiny AC. Diel variability in the elemental composition of the marine cyanobacterium Synechococcus. J Plankton Res. 2016;38: 1052–1061. doi:10.1093/PLANKT/FBV120

39. Roscoff Culture Collection | Marine Microalgae, Macroalgae, Protists, Bacteria and Viruses. [Internet] [cited 7 Apr 2023]. Available from: https://roscoff-culture-collection.org/rcc-strain-details/2673.

40. Roscoff Culture Collection | Marine Microalgae, Macroalgae, Protists, Bacteria and Viruses. [Internet] [cited 7 Apr 2023]. Available from: https://roscoff-culture-collection.org/rcc-strain-details/515.

41. Roscoff Culture Collection | Marine Microalgae, Macroalgae, Protists, Bacteria and Viruses. [Internet] [cited 7 Apr 2023]. Available from: https://roscoff-culture-collection.org/rcc-strain-details/1086.

42. Roscoff Culture Collection | Marine Microalgae, Macroalgae, Protists, Bacteria and Viruses. [Internet] [cited 7 Apr 2023]. Available from: https://roscoff-culture-collection.org/rcc-strain-details/32.

43. Bertilsson S, Berglund O, Karl DM, Chisholm SW. Elemental composition of marine Prochlorococcus and Synechococcus: Implications for the ecological stoichiometry of the sea. Limnol Oceanogr. 2003;48: 1721–1731. doi:10.4319/LO.2003.48.5.1721

44. Ahlgren NA, Rocap G. Diversity and distribution of marine Synechococcus: Multiple gene phylogenies for consensus classification and development of qPCR assays for sensitive measurement of clades in the ocean. Front Microbiol. 2012;3: 213. doi:10.3389/FMICB.2012.00213/BIBTEX

45. Sterner RW, Elser JJ. Ecological Stoichiometry: The Biology of Elements from Molecules to the Biosphere. Princeton University Press; 2002.

46. Grob C, Ulloa O, Claustre H, Huot Y, Alarcón G, Marie D. Contribution of picoplankton to the total particulate organic carbon concentration in the eastern South Pacific. Biogeosciences. 2007;4: 837–852. doi:10.5194/BG-4-837-2007

47. Banse K. Cell volumes, maximal growth rates of unicellular algae and ciliates, and the role of ciliates in the marine pelagial1,2. Limnol Oceanogr. 1982;27: 1059–1071. doi:10.4319/LO.1982.27.6.1059

48. Chisholm SW, Frankel SL, Goericke R, Olson RJ, Palenik B, Waterbury JB, et al. Prochlorococcus marinus nov. gen. nov. sp.: an oxyphototrophic marine prokaryote containing divinyl chlorophyll a and b. Arch Microbiol. 1992;157: 297–300. doi:10.1007/BF00245165/METRICS

49. Webb EA, Ehrenreich IM, Brown SL, Valois FW, Waterbury JB. Phenotypic and genotypic characterization of multiple strains of the diazotrophic cyanobacterium, Crocosphaera watsonii, isolated from the open ocean. Environ Microbiol. 2009;11: 338–348. doi:10.1111/J.1462-2920.2008.01771.X

50. Lomas MW, Baer SE, Acton S, Krause JW. Pumped up by the cold: Elemental quotas and stoichiometry of cold-water diatoms. Front Mar Sci. 2019;6: 286. doi:10.3389/FMARS.2019.00286/BIBTEX

51. Zhang Y, Fu FX, Whereat E, Coyne KJ, Hutchins DA. Bottom-up controls on a mixed-species HAB assemblage: A comparison of sympatric Chattonella subsalsa and Heterosigma akashiwo (Raphidophyceae) isolates from the Delaware Inland Bays, USA. Harmful Algae. 2006;5: 310–320. doi:10.1016/J.HAL.2005.09.001

52. Zinser ER, Johnson ZI, Coe A, Karaca E, Veneziano D, Chisholm SW. Influence of light and temperature on Prochlorococcus ecotype distributions in the Atlantic Ocean. Limnol Oceanogr. 2007;52: 2205–2220. doi:10.4319/LO.2007.52.5.2205

